# Glycogen metabolism in mouse embryonic Sertoli cells sustains the germ line through the lactate shuttle

**DOI:** 10.1101/2025.07.23.666216

**Authors:** Martín A. Estermann, Joseph Sheheen, Sara A. Grimm, Boris Tezak, Yu-Ying Chen, Tsuyoshi Morita, Humphrey H.C. Yao, Blanche Capel

## Abstract

Metabolites are key regulators of cell fate decisions, epigenetic remodeling, and lineage commitment. While genetic pathways governing testis differentiation are well studied, the role of metabolism remains poorly understood. In this study, we investigated a transient, male-specific accumulation of glycogen in supporting (Sertoli) cells of embryonic testis in mice. Although glycogen metabolism was dispensable for Sertoli cell differentiation, its disruption led to reduced lactate production and impaired PGCs ability to colonize the newly forming testis cords. Inhibiting lactate transport further revealed a critical metabolic coupling between Sertoli and germ cells during early testis development. Surprisingly, external lactate or glucose supplementation failed to rescue the germ cell phenotype. These findings suggest that glycogen accumulation supports a critical developmental window in which both Sertoli and germ cells are metabolically constrained and unable to rely on external carbon sources. This highlights a sensitive period during testicular development where nutrient scarcity could have long-term consequences for fertility.

## Introduction

Increasing evidence across developmental systems indicates that metabolism is far more than a background process supporting biosynthesis and energy production. It is an active regulator of cell fate decisions, epigenetic remodeling, and lineage commitment^1–6^. In this context, the concept of metabolic coupling, the coordinated exchange of metabolites between specialized cells, has emerged as a key mechanism by which cell populations communicate, synchronize their developmental trajectories and compartmentalize their functions. This mechanism, often exploited by cancer cells^7–9^, plays a vital role in the normal function of many organs, including in the brain, kidney, muscle and adult testis^10–13^. In the adult testis, Sertoli cells provide metabolic and functional support to germ cells, to sustain sperm production^14–18^. However, when this metabolic coupling is established during testis development has not been explored.

Fetal testicular differentiation involves the transformation of the bipotential gonadal primordium into a testis, a process orchestrated by the supporting cell lineage^19^. Supporting cells are the first to commit to a testicular (Sertoli) fate around embryonic day 11.5 (E11.5) in mice^20–22^. Once specified, Sertoli cells coordinate the differentiation of the rest of gonadal cell types, including the germline^22–24^. While the genetic regulators driving these processes have been well characterized, much less is known about how metabolic cues influence sexual fate decisions.

A puzzling early metabolic difference between XY and XX gonadal supporting cells at the onset of sex determination is the accumulation of glycogen deposits in male, but not female, supporting cells^25,26^. Despite this observation, the exact role of this sex-specific metabolic difference has not been clearly identified. To explore this metabolic dimorphism, we perturbed glycogen synthesis and breakdown pathways. We show that while glycogen metabolism is not required for Sertoli cell differentiation, it is required for germ cells to initially populate the fetal testis cords as they form. Specifically, Sertoli cells break down glycogen to produce lactate, which is then transported to germ cells via the lactate shuttle, supporting their development. These findings establish glycogen metabolism as a critical determinant of germ cell development, highlighting a previously unrecognized metabolic coupling between Sertoli and germ cells during fetal testis development, with potential implications for male fertility.

## Results

### Sertoli cells accumulate glycogen within a specific developmental window

To characterize the glycogen deposition pattern, immunofluorescence was performed using a glycogen-specific antibody^27^ on gonadal paraffin sections from sexually differentiating male and female gonads. No detectable glycogen staining was observed in either male or female gonads at E11.5 (Ts 20) (Fig. S1A). Despite the presence of SRY+ and SOX9+ Sertoli cells in E11.5 testes, glycogen was not detected at this developmental stage, suggesting that glycogen accumulates after sex differentiation (Fig. S1B). As a positive control, the notochord displayed strong glycogen immunoreactivity at E11.5 (Fig. S1C). By E12.5, glycogen was detected in Sertoli cells in the testicular cords, colocalizing with a known Sertoli cell marker AMH, but was absent in FOXL2 positive ovarian supporting (pre-granulosa) cells (Fig 1A). Interestingly, by E13.5, glycogen was no longer detectable in the testis (Fig. 1A). This suggests a tightly regulated developmental window in which glycogen accumulates in Sertoli cells between E11.5 and E12.5, followed by its breakdown and/or consumption between E12.5 and E13.5.

**Fig. 1:**
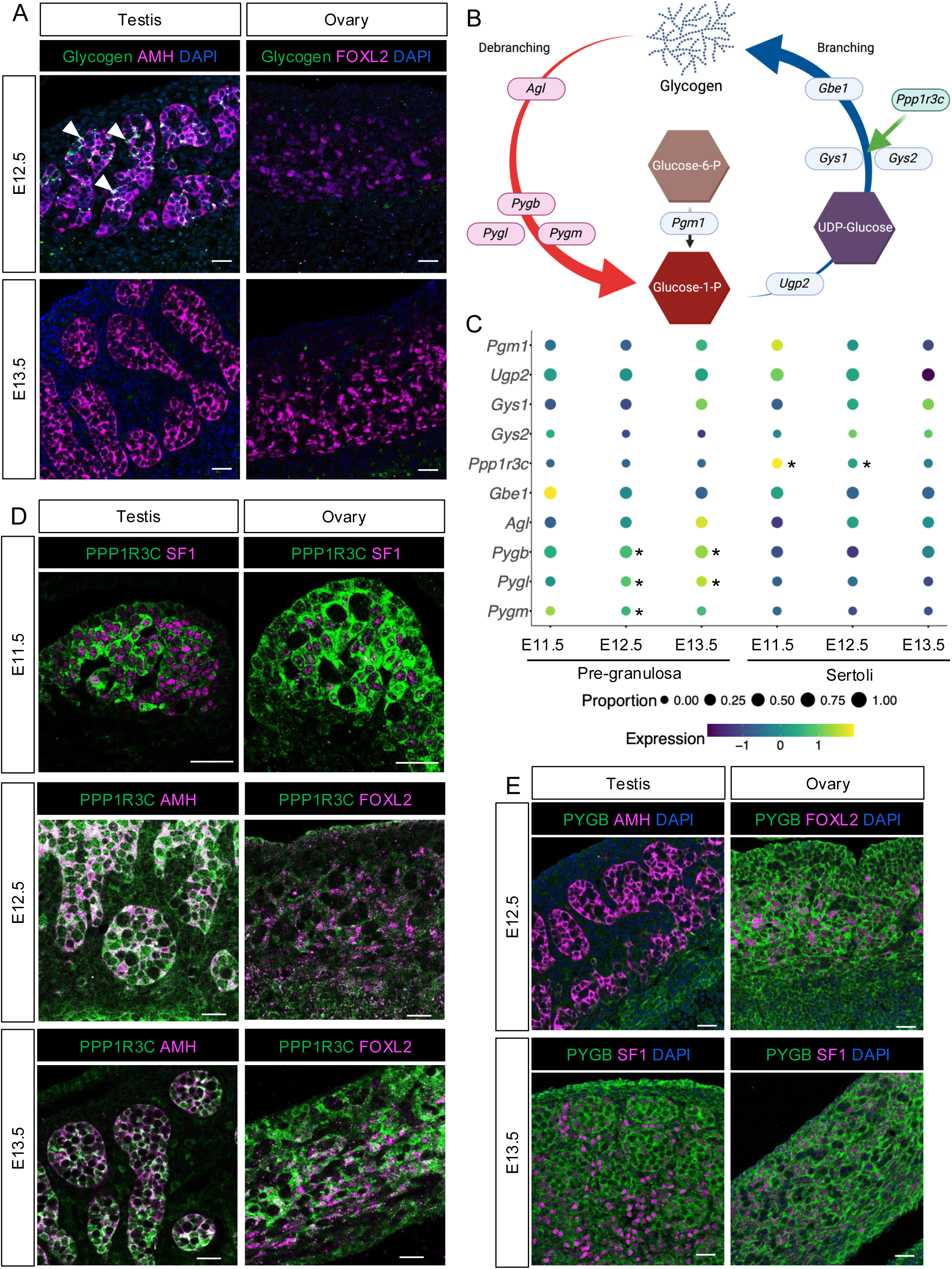
Differential glycogen metabolism results in Sertoli cell-specific accumulation. (A) Immunofluorescence for glycogen (green) and male (AMH) or female (FOXL2) supporting cell markers (magenta) in E12.5 and E13.5 gonads. White arrowheads indicate Glycogen deposition in Sertoli cells. Samples were counterstained with DAPI (blue). (B) Schematic representation of the glycogen synthesis (right) and glycogen degradation (left) pathway (created with BioRender.com). (C) Dot plot representation of the expression of glycogen pathway related genes in male (right) and female (left) supporting cells at different developmental timepoints. * indicate adjpvalue<0.05, Wilcoxon Rank Sum test. (D) Immunofluorescence for PPP1R3C (green) and supporting cell markers SF1, AMH or FOXL2 (magenta) in transverse (E11.5) or longitudinal (E12.5 and E13.5) gonadal sections. (E) Immunofluorescence for PYGB (green) and supporting cell markers SF1, AMH or FOXL2 (magenta) in E12.5 and E13.5 gonads. Samples were counterstained with DAPI (blue). Scale bars are 50 μm.

### Sex-specific glycogen synthesis and degradation result in glycogen accumulation in Sertoli cells

To understand the molecular mechanism underlying glycogen accumulation and subsequent breakdown, we interrogated a mouse gonadal single-nucleus RNA-seq dataset^28^ for the expression of enzymes involved in glycogen synthesis and breakdown in supporting cells across developmental stages (Fig. 1B-C). No significant differences were observed in the expression of glycogen synthesis enzymes (*Pgm1*, *Ugp2*, *Gys1*, *Gys2* and *Gbe1*) (Fig. 1C). However, Sertoli cells in the embryonic testis expressed significantly higher levels of *Ppp1r3c* at E11.5 and E12.5 than the pre-granulosa cells of embryonic ovary (Fig. 1C, Table S1). *Ppp1r3c*, also known as protein targeting to glycogen (PTG), regulates the phosphorylation (activation) of glycogen synthase (GSY1 and GSY2), therefore promoting glycogen formation. At E12.5, Sertoli cells showed higher PPP1R3C protein expression than pre-granulosa cells, while expression levels were comparable at E11.5 and E13.5 (Fig. 1D). This temporal difference in expression aligns with the pattern of glycogen accumulation observed in the testis (Fig. 1A).

In contrast, pre-granulosa cells exhibited significantly higher levels of enzymes that degrade glycogen, *Pygl*, *Pygb* and *Pygm* at E12.5 and *Pygl* and *Pygb* at E13.5 (Fig. 1C, Table S1). PYGB, the most abundant isoform (Fig. 1C, Table S1), was expressed at similar levels in XX and XY gonads at E11.5 and E13.5 (Fig. S1D, 1E). However, testes exhibited lower expression of PYGB at E12.5 (Fig. 1E). These data suggest that glycogen accumulates in Sertoli cells at E12.5 as the result of differential synthesis (mediated by PPP1R3C) and reduced glycogen breakdown (mediated by PYG enzymes).

### Glycogen is not required for fate determination of Sertoli cells

To investigate the role of glycogen breakdown during gonadal differentiation, we performed *ex-vivo* gonadal cultures using the glycogen phosphorylase inhibitor (GPI) CP 316819, which inhibits all 3 glycogen phosphorylase enzymes (PYGB, PYGL and PYGM). Taking advantage of the gonads being paired organs, we incubated one E12.5 gonad from a single embryo with the GPI inhibitor and the contralateral gonad with a vehicle control for 48 hours. To determine the optimal concentration and assess potential toxicity, we cultured gonadal pairs with 75 µM, 100 µM and 150 µM of GPI or vehicle solutions and evaluated apoptosis via cleaved caspase 3 (CC3) staining and proliferation by staining for KI-67 (MKI-67) (Fig. S2A). We selected 75 µM of GPI for further experiments, as higher concentrations increased gonadal apoptosis (Fig. S2A). We first examined whether glycogen breakdown was required for Sertoli cell differentiation. Although gonads grew more slowly in vitro and did not retain perfect morphology, male gonads cultured with 75 µM of GPI exhibited normal Sertoli cell differentiation, evidenced by the expression of SOX9 and the absence of FOXL2, a marker of pre-granulosa cells (female vehicle control) as indicative of sex reversal (Fig. 2A). These findings suggest that glycogen breakdown is not required for Sertoli cell differentiation or fate maintenance.

**Fig. 2:**
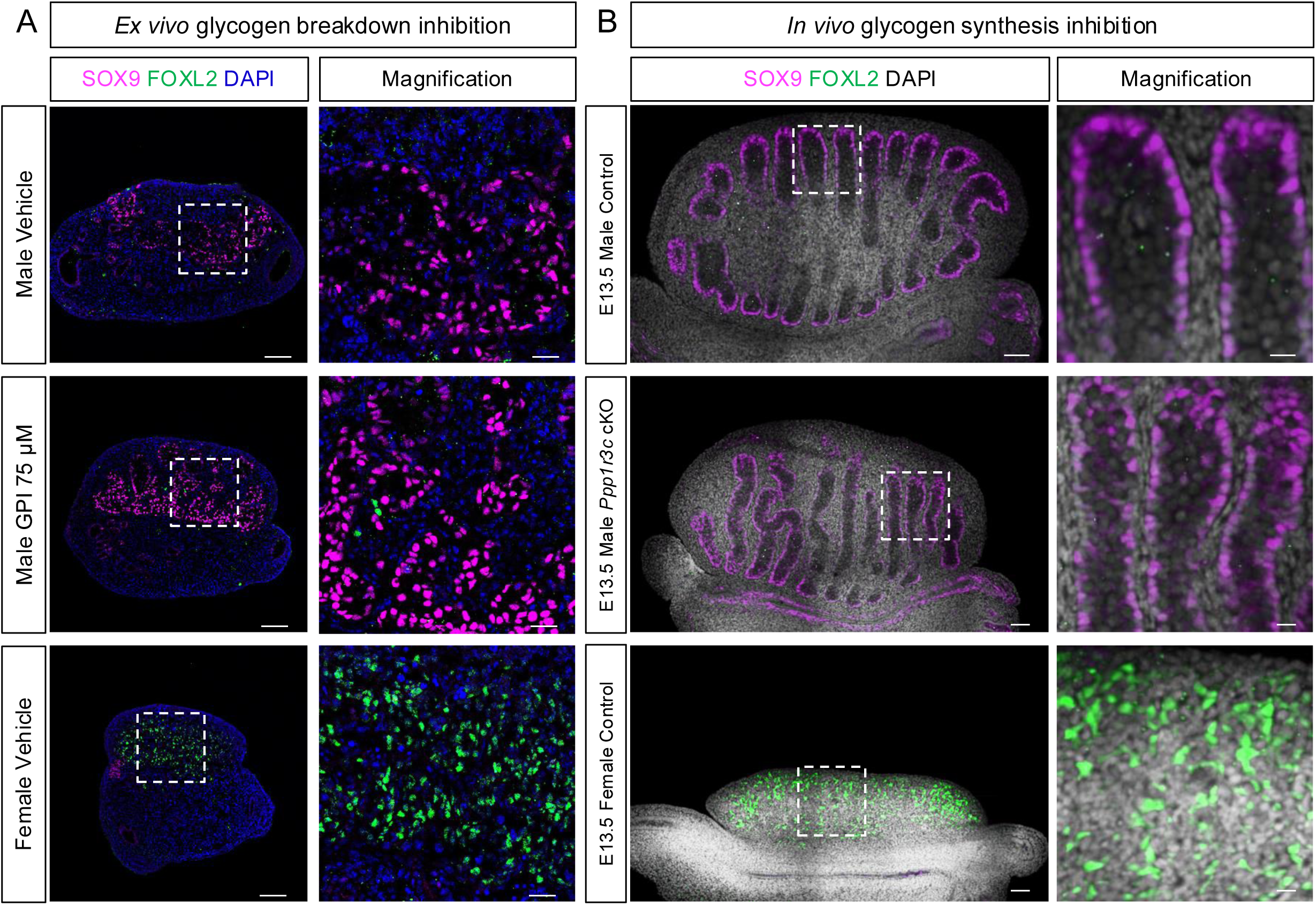
Glycogen is not required for Sertoli cell differentiation. (A) Immunofluorescence for the Sertoli cell marker SOX9 (magenta) and pre-granulosa cell marker FOXL2 (green) in cultured gonads treated with 75 µM GPI or vehicle solution. Samples were counterstained with DAPI (blue) (scale bars=100 μm). White dashed box indicates the magnified area (scale bars=25 μm. (B) Immunofluorescence for the Sertoli cell marker SOX9 (magenta) and pre-granulosa cell marker FOXL2 (green) in E13.5 male and female control and male *Ppp1r3c* cKO gonads. Samples were counterstained with DAPI (grey) (scale bars=50 μm). White dashed box indicates the magnified area (scale bars=12.5 μm).

To evaluate the role of glycogen synthesis in Sertoli cells, we obtained a *Ppp1r3c* conditional mouse line^29^ and generated *Ppp1r3c* conditional knockout (cKO) mice using *Nr5a1*-Cre, a Cre expressed in progenitors for Sertoli cells. To confirm that deletion of *Ppp1r3c* leads to the abrogation of glycogen accumulation in Sertoli cells, we performed periodic acid-Schiff (PAS) staining, which detects the presence of polysaccharides, including glycogen, on E12.5 control or *Ppp1r3c* cKO gonads (Fig. S2B). In control gonads, as expected, PAS positive staining was detected in Sertoli cells (Fig. S2B), consistent with our previous results (Fig. 1A). In contrast, PAS staining was absent in cKO testes, similar to results in the ovary, (Fig. S2B), confirming that loss of *Ppp1r3c* blocks glycogen synthesis and its accumulation. Immunofluorescence for Sertoli (SOX9) and pre-granulosa (FOXL2) cell markers indicated that loss of *Ppp1r3c* in Sertoli cells did not affect Sertoli cell differentiation (Fig. 2B). These results collectively demonstrate that neither the synthesis nor the degradation of glycogen is required for Sertoli cell fate determination.

### Altered glycogen metabolism in Sertoli cells results in germ cell delay

To investigate the role of glycogen breakdown in gonadal differentiation, we performed bulk RNA-seq on paired testes cultured with 75 µM of GPI or vehicle solution. A total of 5 testicular pairs were sequenced, and principal component analysis (PCA) revealed a clear separation by treatment (Fig. S3A). Paired differential expression analysis identified 3,202 differentially expressed genes (DEGs), including 1,770 downregulated and 1,432 upregulated in the GPI-treated testes compared to controls (Fig. S3B, Table S2). Pathway analysis of the upregulated genes identified lipid metabolism among the most enriched pathways (Fig. 3A, Table S2), suggesting a metabolic shift from glucose oxidation to fatty acid metabolism. Pathway analysis of the downregulated genes identified an impairment of genes involved in extracellular matrix organization and in germ cell development (Fig. 3A). To determine whether these transcriptional changes in germ cell gene expression reflected changes in germ cell numbers, we performed whole mount immunofluorescence for the pan-germ cell marker DDX4 (Fig. 3B) and quantified the germ cell numbers in control and GPI treated testes (Fig. 3B-C). GPI-treated testes exhibited a significant reduction in germ cell number, with an average of 91% fewer germ cells, compared with the control (Fig. 3B-C), suggesting that glycogen degradation in Sertoli cells is essential for germ cell population of the gonad.

**Fig. 3:**
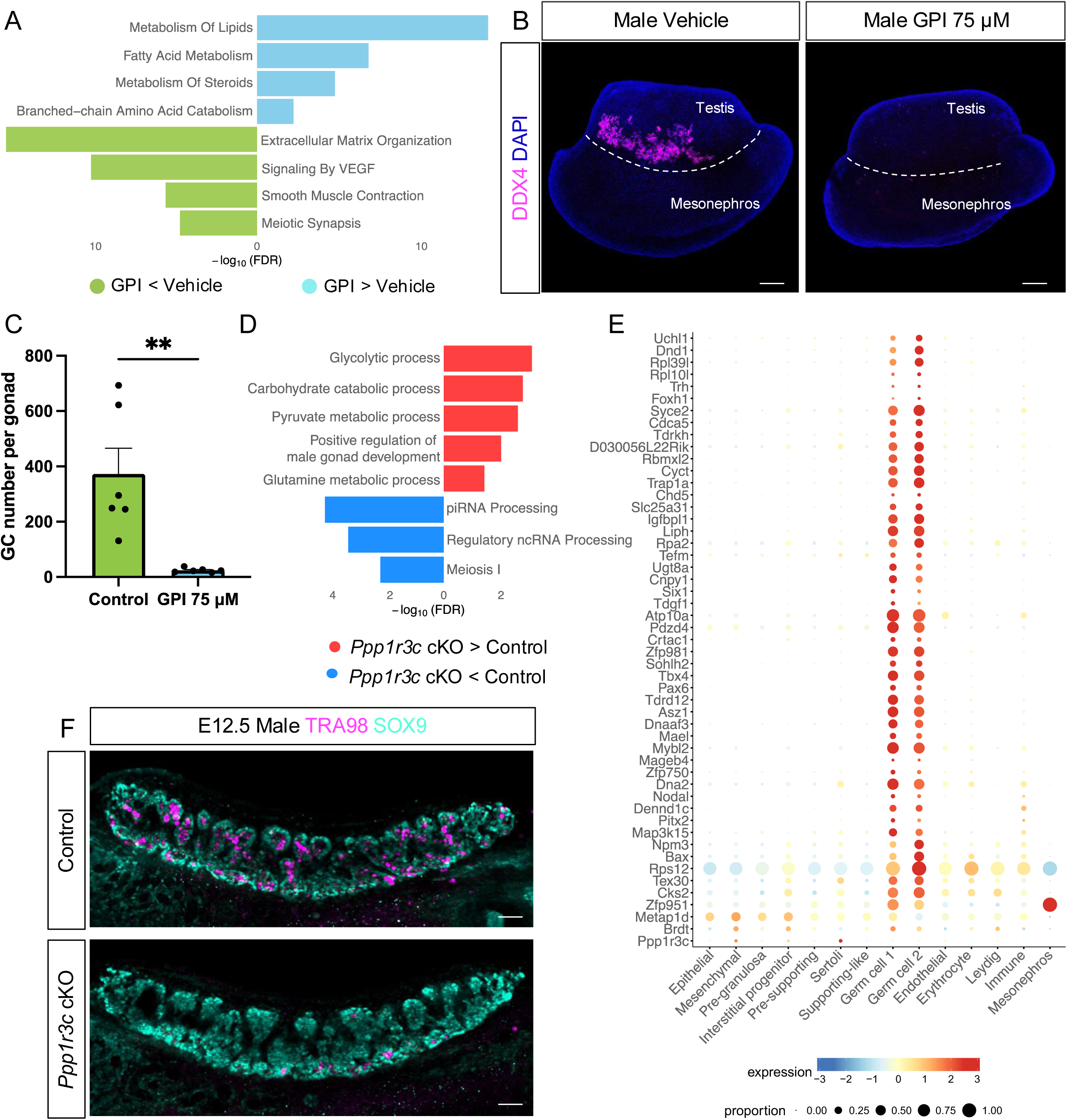
Blocking glycogen breakdown or synthesis reduces germ cell number in XY gonads during sex determination. (A) Reactome pathway analysis of significant upregulated (cyan) and downregulated (green) genes in the GPI treated gonads vs the vehicle control. (B) 3D projection of DDX4 (germ cell, magenta) whole mount immunostaining in male gonads cultured with 75 µM GPI or vehicle solution. Samples were counterstained with DAPI (blue). Dashed white line delineates the testicular region. Scale bars=100 μm. (C) Quantification of the total number of germ cells in male gonads cultured with 75 µM GPI or vehicle solution. Bars represent mean±s.e.m., n=6. Two-tailed t-test. **p<0.01. (D) Gene ontology (BP) analysis of significant upregulated (red) and downregulated (blue) genes in E12.5 *Ppp1r3c cKO* testis vs control. (E) Dot plot representation of the expression of all the downregulated genes in the *Ppp1r3c* cKO testes in different embryonic E12.5 testicular cell populations from the single-nucleus RNA-seq dataset. (F) Immunofluorescence for the Sertoli cell marker SOX9 (green) and germ cell marker TRA98 (magenta) in E12.5 male control and *Ppp1r3c* cKO gonads. Scale bars=50 μm.

As the GPI treatment targets all gonadal cells, the role of Sertoli specific glycogen metabolism in gonadal differentiation was evaluated using our mouse cKO model. Bulk RNA-seq in control and *Ppp1r3c* cKO E12.5 testis (Fig. S3C) and differential expression analysis identified 111 differentially expressed genes, 57 upregulated and 54 downregulated (Fig. S3D, Table S3). Notably, *Ppp1r3c* was the top downregulated gene, with a 76.8% reduction, confirming a robust gene knockout. Pathway analysis of the differentially upregulated genes revealed enrichment in carbohydrate metabolism pathways, suggesting either a compensatory metabolic response to glycogen loss or possibly more available glucose in the absence of glycogen reservoirs (Figure 3D, Table S3). Interestingly, and consistent with our inhibitor results, downregulated genes were associated with germ cell functions, including piRNA processing and meiosis (Fig. 3D). Notably, almost all (50/54) downregulated genes were exclusively expressed in germ cells (Fig. 3E), indicating a direct impact on the germline. To determine whether these transcriptional changes in germ cell gene expression reflected changes in germ cell numbers, we performed whole mount immunofluorescence for the germ cell marker TRA98 (Fig. 3E, S3E-F) and quantified germ cell numbers in control and *Ppp1r3c* cKO testes (Fig. S3G). While germ cell numbers were comparable between control and cKO gonads at E11.5 (Fig. S3E, S3G), by E12.5 a significant reduction of germ cell number was observed in *Ppp1r3c* cKO testes (Fig. 3F, S3G). However, by E13.5 as glycogen deposition declined in Sertoli cells, germ cell numbers recovered (Fig. S3F-G). Taken together, these findings suggest embryonic Sertoli cell glycogen metabolism is utilized by germ cells to populate the fetal testis during a transient window of development.

### Sertoli and germ cells are metabolically coupled through the lactate shuttle

As their name suggests, supporting cells play a crucial role in nourishing the germline, raising the possibility that Sertoli cells and germ cells are metabolically coupled. Because lactate is a common product of glycogen breakdown^30^ and because meiotic and post-meiotic germ cells in the adult testis rely on lactate provided by Sertoli cells^14–18^, we investigated whether this system might underlie the effects of glycogen loss on germ cell numbers.

As our findings suggest that Sertoli cells break down their stored glycogen to fuel the germline, we hypothesized that glycogen is oxidized into lactate. Indeed, both inhibition of glycogen breakdown using GPI and inhibition of glycogen synthesis in the *Ppp1r3c* cKO model resulted in an approximately 40% reduction in intracellular lactate levels in embryonic testes (Fig. 4A-B), strongly suggesting that lactate is a significant by-product of glycogen breakdown in early Sertoli cells.

**Fig. 4:**
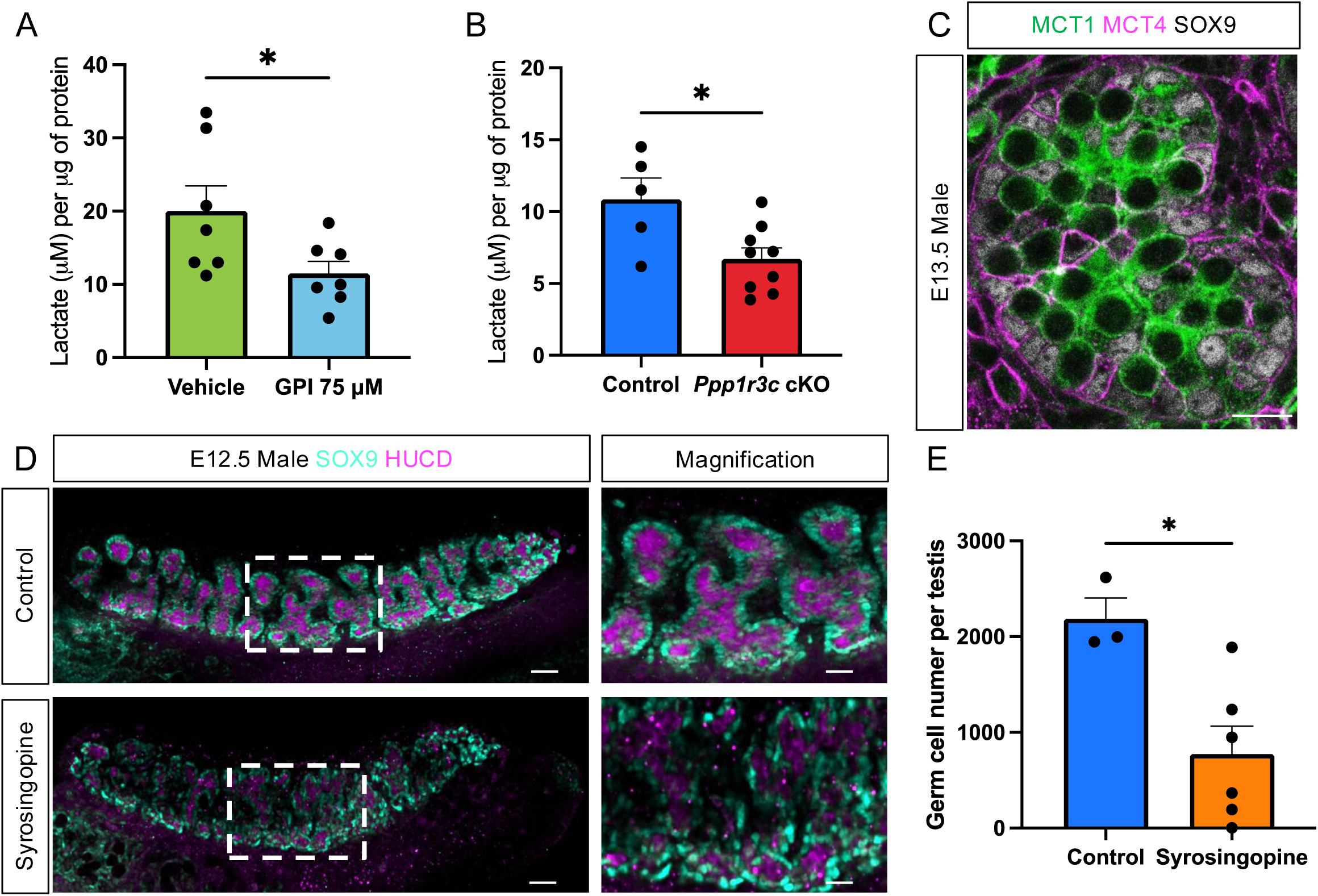
Glycogen metabolism in Sertoli cells sustains germ cell survival through lactate shuttling. (A) Lactate concentration in testis cultured with 75 µM GPI or vehicle solution. Bars represent mean±s.e.m., n=7. Two-tailed t-test. *p<0.05. (B) Lactate concentration in E12.5 *Ppp1r3c cKO* or control testes. Bars represent mean±s.e.m., n≥5. Two-tailed t-test. *p<0.05. (C) Immunofluorescence for MCT1 (green), MCT4 (magenta) and SOX9 (grey) in E13.5 male gonads. Scale bar=12 μm. (D) Immunofluorescence for the Sertoli cell marker SOX9 (cyan) and germ cell marker HUCD (magenta) in E12.5 male control and Syrosingopine treated gonads (scale bars=50 μm). White dashed box indicates the magnified area (scale bars=25 μm). (E) Quantification of the total number of germ cells per testis from control and Syrosingopine treated mice. Bars represent mean±s.e.m., n≥3. Two-tailed t-test. *p<0.05.

Lactate moves between cells through the exporter MCT4 (*Slc16a3)* and the importer MCT1 (*Slc16a1)*, also known as the lactate shuttle. To determine whether lactate produced in Sertoli cells could be transported into the germ cells through this shuttle, we examined the presence of MCT4 and MCT1 proteins by immunofluorescence and found that the lactate exporter (MCT4) was predominantly present in SOX9-positive Sertoli cells whereas the importer (MCT1) was found in germ cells (Fig. 4C). These data indicate that the machinery to transport lactate from Sertoli to germ cells is in place and support the notion of metabolic coupling between these cell populations during embryonic development.

To investigate whether this metabolic coupling is required for germ cell survival, we injected a dual MCT1 and MCT4 inhibitor (Syrosingopine)^31^ or vehicle solution into pregnant female mice at the onset of embryonic sex determination (E10.5 and E11.5) and examined the testes from the XY embryos at the height of glycogen deposition (E12.5) (Fig. 4D). Syrosingopine treatment in vivo did not affect Sertoli cell fate; however, germ cell numbers were significantly decreased (Fig. 4D-E). This result phenocopied the glycogen depletion results (Fig. 3), consistent with the model whereby glycogen is metabolized into lactate in Sertoli cells, then transported to germ cells through the lactate shuttle during this pivotal window of testis differentiation.

### The decreased germ cell numbers due to inhibition of glycogen breakdown cannot be rescued by providing exogenous lactate or glucose

Based on the results that inhibition of glycogen degradation resulted in lower levels of lactate (Fig. 4A-B), we wondered if exogenous lactate could rescue germ cell loss in GPI treated gonads. To test this hypothesis, we cultured embryonic testes with either GPI (75 µM), or GPI (75 µM) plus sodium lactate (25 mM) (Fig. S4A). Consistent with our previous results, treatment with GPI alone led to a reduction of germ cell number (Fig. S4A-B, S3E). Contrary to our expectations, supplementation of GPI-treated gonads with an exogenous supply of lactate (25mM sodium lactate) failed to rescue the germ cell phenotype (Fig. S4A-B, S4E), suggesting that lactate cannot fully compensate for the inhibition of glycogen breakdown. One possible explanation is that germ cells, which are closely surrounded by Sertoli cells and enclosed in the center of testis cords, do not have direct access to extracellular lactate. Because Sertoli cells do not express the lactate importer, they cannot act as intermediaries in its uptake and distribution. To evaluate if glucose could be used as a source for lactate generation in Sertoli cells, bypassing glycogen degradation, we cultured the embryonic testes with either GPI or GPI with 25 mM of D-glucose (Fig. S4C-E). Despite a small, but non-significant, increase in germ cell number (Fig. S4D), glucose supplementation, similar to lactate supplementation, failed to lead to normal germ cell numbers (Fig. S4C-E).

Overall, these findings suggest that Sertoli cells accumulate glycogen and break it down within a transient developmental window to support germ cell survival, as neither Sertoli cells nor germ cells can rely on external sources of carbons like glucose and lactate during this period. This reveals a metabolic pairing during a sensitive period of testicular development which has long-term consequences on fertility^32–34^.

## Discussion

As glycogen deposition is one of the earliest recognized sexual dimorphisms during mouse gonadal sex differentiation^25^, it has been an open question whether glycogen directly impacts the differentiation process of the male gonad. Here we found that neither glycogen synthesis nor its breakdown is required for SOX9 expression, Sertoli cell differentiation or testis morphological patterning. This was unexpected, given previous findings that high extracellular glucose is essential for SOX9 expression and testicular differentiation^26^, suggesting that glucose derived from glycogen stores and glucose acquired through uptake may have different metabolic fates during early gonadal development.

Disrupting glycogen metabolism in embryonic Sertoli cells, through an organ culture or genetic approach, significantly reduced the number of testicular germ cells. We investigated whether this phenotype could be explained by loss of lactate, a common product of glycogen breakdown and a known nutrient for adult spermatogonia^14,30^. We found that levels of lactate are reduced when either deposition or breakdown of glycogen is blocked. Furthermore, blocking lactate intercellular transport in vivo phenocopies the germ cell loss phenotype we observed when glycogen was depleted. These findings support the model that lactate derived from compartmentalized glycogen in Sertoli cells is an essential metabolite for germ cells entering the fetal testis cords

Matoba et al. were the first to identify and characterize the sex-specific glycogen accumulation in the developing testis using PAS staining^25^. PAS-positive signal was first detected between E11 and E11.5 (14ts to 18ts), coinciding with the expression of SOX9 in Sertoli cells^25^. Interestingly, when we used a glycogen-specific antibody^27^ we were unable to detect glycogen signal in SOX9-positive E11.5 testis, despite robust staining in the notochord. These differences could be attributed to different factors, including strain-specific developmental timing differences (outbred ICR vs inbred C57BL/6)^35^ or differences in sensitivity and specificity between the two glycogen detection methods. While PAS staining lacks specificity, detecting a range of polysaccharides including glycogen, it can identify glycogen precursors such as unbranched glucose chains, which may not be recognized by the antibody. Consequently, while PAS staining is likely more sensitive for detecting early or precursor forms of glycogen, antibody-based detection offers greater specificity for mature glycogen, highlighting the value of combining both approaches to capture the dynamics of glycogen accumulation during early testicular development.

Two orthogonal methods, the in vitro chemical inhibition of glycogen degradation (GPI inhibitor) and the in vivo conditional knockout of *Ppp1r3c* (an enzyme required for glycogen deposition), were used here to interrogate the role of glycogen during sex determination. Although these two approaches examined the same questions from different angles, together they provide a broader understanding of the function of stored glycogen during testis differentiation. For example, stopping glycogen breakdown may trap essential glycolytic metabolites while allowing normal function of glycogen branching machinery. Conversely, stopping glycogen buildup impairs normal glycogen branching machinery but does not lock limited glucose into glycogen deposits. By combining these two methods we can be more confident that the phenotypes are not secondary to the reduced systemic metabolic input or altered metabolic signaling.

Metabolic coupling similar to the one we characterized between fetal Sertoli and germ cells is also observed in other systems. In the central nervous system, astrocytes, which are essential for the blood-brain barrier formation and maintenance, take up glucose from capillaries to replenish their glycogen storage, which can be rapidly turned over to lactate and exported through the lactate shuttle when neurons require energy^30,36–39^. Similarly, postnatal Sertoli cells establish the blood-testis barrier and appear to stockpile glycogen and convert it to lactate as a source of energy for germ cells^40–46^. However, a role for metabolic coupling between Sertoli cells and early germ cells in the fetal testis had not been explored previously. We discovered that metabolic interplay between supporting cells and germ cells occurs as soon as germ cells enter the fetal testis. Interestingly, this developmental period coincides with the differentiation window when male germ cells switch from glycolytic metabolism to oxidative phosphorylation^47,48^. It is likely that lactate derived from glycogen degradation in Sertoli cells is a key enabler of this switch in vivo as lactate allows an expedited metabolic path to oxidative phosphorylation without the energetic requirements of glycolysis. Directly testing this will be necessary to determine the full mechanism of Sertoli cell directed male germ cell metabolic reprograming. It is unclear if germ cell recovery is due to successful completion of a metabolic switch in germ cells or enhanced availability of nutrients once Sertoli cells have completed cord formation.

Chemical inhibition of glycogen degradation led to a significant reduction in available intracellular lactate, impairing the ability of cells to meet germ cell demands. Given this deficit, it was expected that exogenous supplementation with glucose (product of glycogen degradation) or lactate would bypass the need for glycogen-derived energy^30,49^. Surprisingly, supplementation with either lactate or glucose failed to rescue the germ loss phenotype. These findings suggests that glycogen accumulation is essential to support a critical developmental window during which neither Sertoli nor germ cells can rely on external carbon sources. Notably, the developing testis undergoes major structural changes during the period of active glycogen metabolism (E11.5 to E13.5)^19^, including an extensive vascular remodeling and testis cord formation^50,51^. The morphological reorganization of the XY gonad likely imposes high energy demands, prompting Sertoli cells to use compensatory mechanisms, such as glycogen storage and compartmentalization, to simultaneously support the metabolic needs of developing male germ cells. Male germ cells clonally proliferate and die during this temporally constrained period yielding a selected cohort that insures fertility in adulthood^32,33^. Thus, disrupting the timing of proliferative waves likely has lasting impacts on stemness and fertility. Although, the *Ppp1r3c* cKO mice have a ∼50% incidence of infertility and Sertoli only seminiferous cords in adulthood (unpublished data), whether or not this is caused by fetal defects cannot be determined in our system.

Together these results provide a compelling example of the emerging field of metabolofertility^52^, highlighting a critical window of metabolic sensitivity during which male germ cells depend on lactate derived from Sertoli cell glycogen stores to populate the newly forming testis cords. This work adds to a growing body of evidence linking metabolism and fertility, demonstrating how metabolic status, pathways, and intercellular metabolite exchange influence reproductive function across multiple levels, including gametogenesis.

## Materials and Methods

### Animals

C57BL/6 mice were purchased from Jackson Laboratory (stock number 000664). *Nr5a1-Cre* mice (B6D2-g(Nr5a1-cre)2Klp) were generated by Keith Parker and maintained on a C57BL/6 background^53^. *Ppp1r3c* floxed mice^29^ were obtained from the Saltiel Lab at UCSD and maintained on a C57BL/6 background.

Pairs were set up overnight and females were screened for the presence of vaginal plug the next morning (E0.5). Embryos were staged by tail somite number upon embryonic harvest. For genotyping, DNA was isolated from individual embryos by overnight tissue dissociation in proteinase K, followed by 70% isopropyl alcohol precipitation. Genotyping was performed by PCR and subsequent agarose gel electrophoresis. (*Ppp1r3c*: CCTTTATAGTTGGACCTGTCATGG, TCTACAGTCTAGCTCTGTGCTTGG, expected band size: 656bp wildtype *Ppp1r3c*, 741bp floxed *Ppp1r3c*; *Nr5a1-Cre*: GAACCTGATGGACATGTTCAGG, AGTGCGTTCGAACGCTAGAGCCTGT, expected band size: 320bp; Sex genotyping: TCATGTCCATCAGGTGATGG, CAATGTGGACCATGACATTG, ATGGACACAGACATTGATGG, expected band size: 420bp *utx*, 288 bp *uty*).

All animal procedures at the National Institute of Environmental Health Sciences were approved by NIEHS Animal Care and Use Committee and are in compliance with a NIEHS-approved animal study proposal (2010-0016). At Duke University Medical Center, all mice were housed in accordance with National Institutes of Health guidelines and the approval of the Duke University Medical Center Institutional Animal Care and Use Committee (A089-20-04 9N).

### GPI gonadal cultures

Individual gonadal cultures were performed using a modified version of the agar gonad culture method^54^. Agar slabs suitable for 48-well plates were generated using newly designed 3D-printed molds (Fig. S4F). Briefly, 150 mg of BactoAgar (214010, Difco Labs) was dissolved in 10 ml of complete media, consisting of 10% heat-inactivated fetal bovine serum (16140071, Gibco), 1X Penicillin-Streptomycin (P0781, Sigma-Aldrich) in 1X phenol red-free DMEM/F12 (1:1) (21041-025, Gibco). The solution was microwaved in 5 seconds intervals until fully dissolved. Then, 90 µl of the agar solution was added to each mold and solidified agar slabs were transferred to a 48 well plate.

Individual paired gonad-mesonephros complexes were isolated from E12.5 C57BL/6 mouse embryos in cold PBS. Gonads were transferred to the agar slabs containing 180 µl of complete media supplemented with either CP 316819 inhibitor (GPI) (PZ0189, Sigma-Aldrich) at the desired concentration or vehicle solution (DMSO). Gonads were cultured for 48 hours at 37°C in 5% CO2, replacing 100 µl of fresh media with an inhibitor or vehicle every 24 hours. Cultured gonad-mesonephros complexes were imaged in a Leica MZ16 Stereo Microscope with a Leica MC190 HD Microscope Camera. For rescue experiments, media was supplemented with 25 mM of Sodium L-Lactate (L7022, Sigma-Aldrich) or 25 mM of D-glucose (4912-12, Macron Chemicals).

### Immunofluorescence and histological analyses

C57BL/6 fetal gonad–mesonephros complexes were fixed in 4% PFA overnight at 4 °C. Samples were then washed in PBS, stored at 4 °C in 70% ethanol and paraffin embedded. 5 µm paraffin sections were dewaxed and rehydrated. Cultured gonads were collected and fixed in 4% PFA overnight at 4 °C. Samples were washed in PBS and cryo-protected in 30% sucrose, blocked in OCT and snap frozen. 10 µm frozen sections were cut. Paraffin sections and cryosections were subjected to citrate-based antigen retrieval (Vector Labs, H-3300-250). Samples were transferred to the Sequenza manual immunohistochemistry system, blocked and permeabilized in 0.1% triton X-100 in 1X PBS with 5% normal donkey serum for 1h at RT. Slides were incubated with primary antibodies (Table S4) diluted in blocking buffer at 4 °C overnight. Samples were then washed and incubated with secondary antibodies (Table S4) diluted in blocking buffer at room temperature for 1h. Slides were washed and autofluorescence was quenched using the TrueView Autofluorescence Quenching Kit (Vector Labs, SP-8400). Samples were counterstained with DAPI (Invitrogen, D1306) and mounted using ProLong™ Diamond Antifade Mountant (Invitrogen, P36970). Imaging was performed on a Zeiss LSM 900 confocal microscope using Zen software. Brightness and contrast of images were adjusted using FIJI^55^.

For *Ppp1r3c* mice, gonad–mesonephros complexes were dissected in PBS and fixed in 4% PFA for 30 minutes rocking at room temperature, then dehydrated in an increasing methanol series for long-term storage at −20 °C. Samples for cryo-sectioning were gradually rehydrated, cryoprotected through a sucrose gradient (10%, 15%, 20%, and 30%), embedded in OCT and moved to −80 °C for a minimum of 12 hours before cryo-sectioning. Blocks were serially sectioned at 10-16 μm and placed on slides that were stored at −20 °C until ready to use. For immunostaining, sections were rehydrated in PBS, permeabilized in PBST (0.1% Triton X-100), and blocked for 1 h at room temperature (PBS 0.1% TritonX-100, 3% BSA, 10% horse serum). Sections were incubated in primary antibodies (Table S4) diluted in blocking solution at 4 °C overnight. Following three 10-minute washes in PBS 0.1% TritonX-100, sections were incubated for 2 h at room temperature with DAPI and secondary antibodies (Table S4) diluted in blocking solution. Following three 10-minute washes in PBST, slides were mounted for imaging with polyvinyl alcohol mounting solution and stored at 4 °C. Sections were imaged using the Andor Dragonfly Spinning Disk confocal microscope.

### Periodic Acid Schiff (PAS) staining

The PAS staining protocol was adapted for use on cryo-sectioned tissue. Briefly 10-16 um sections were transferred to superfrost PLUS slides and stored at −20°C until use. Using a pap pen to contain liquid, a 0.5% Periodic acid solution was added to each slide for 5 min at room temperature. Each slide was rinsed 3 times in dH_2_0 and Schiff’s solution was added and incubated for 15 min at room temperature. Each slide was rinsed 3 times in lukewarm tap water and dehydrated in a 25%, 65%, and 100% ethanol gradient. All sectioned samples were mounted in Polyvinyl Alcohol mounting media using #1.5 coverslips and sealed with nail polish. Stained sections were brightfield imaged using an Zeiss axio imager upright microscope.

### Whole mount immunofluorescence

Cultured gonads were collected, fixed in 4:1 methanol:DMSO and stored at −20 °C for at least 24hs. Samples were washed with 50% methanol diluted in 1X PBS for 30 minutes at room temperature, followed by 3 one-hour washes with blocking solution (10% donkey serum and 1% triton in 1X PBS) at room temperature. Samples were incubated overnight at 4°C in blocking solution containing primary antibodies (Table S4). After primary incubation, samples were washed 3 times with blocking buffer for 1 hour each at 4°C and incubated with secondary antibodies (Table S4) in blocking solution overnight at 4°C. Samples were dehydrated by 1-hour washes in increasing methanol concentrations (25%, 50%, 75% with DAPI, and 100%). Tissues were cleared with 1:2 benzyl alcohol:benzyl benzoate (BABB) for at least 24 hours and imaged on a Zeiss LSM 900 confocal microscope using Zen software.

For *Ppp1r3c* mice, gonad–mesonephros complexes were dissected in PBS and fixed in 4% PFA for 30 min rocking at room temperature, then taken up a Methanol gradient to 100% MeOH for long-term storage at −20 °C. After gradual rehydration into PBS, tissues were permeabilized in PBS 0.1 to 0.5% TritonX-100 for 1 h at room temperature and transferred into blocking solution (PBS 0.1 to 1% TritonX-100, 3% BSA, 10% horse serum) for 2 h. Samples were incubated with primary antibodies (Table S4) diluted in blocking solution overnight at 4 °C. Following three 30-minute washes in PBS 0.1 to 1% TritonX-100, tissues were incubated with secondary antibodies (Table S4) and DAPI diluted in blocking solution overnight at 4° C. Tissues were washed three times in PBS/TritonX-100 for 1 h at room temperature, mounted for imaging in 2% Agarose in Cubic R2 clearing media^56^ and stored at 4 °C until imaging on the Andor Dragonfly Spinning Disk confocal microscope. For gonad samples ranging from E11.5-E12.5, the concentration of Triton X-100 was 0.5%, while for samples from E13.5 and above, the concentration was increased to 1%.

### RNA extraction

Vehicle and GPI treated gonads were manually dissected from the mesonephros and RNA was extracted from each vehicle and GPI treated gonadal pair for a total of 5 paired samples. Two E12.5 gonadal pairs (control or Ppp1r3c cKO) were pooled per sample, for a total of 3 samples per group. RNA was extracted using the Arcturus™ PicoPure™ RNA Isolation Kit (Applied Biosystems, KIT0204). Samples were dissociated in 100 µl of extraction buffer with a microtube homogenizer and incubated at 42°C for 30 minutes, then frozen at - 80°C for at least one hour. RNA was precipitated with 100 µl of 70% ethanol and loaded into a pre-conditioned column. Samples were subjected to DNA removal using the RNase-Free DNase Set (Qiagen, #79254). Samples were eluted in 12 µl of elution buffer. RNA was quantified using Qubit™ RNA High Sensitivity kit (Q32852, ThermoFisher Scientific). RNA quality was assessed using High Sensitivity RNA or RNA ScreenTape assay, in a TapeStation (Agilent).

### RNA sequencing

For paired vehicle and GPI treated samples, libraries were made using Tecan’s Ovation® RNA-Seq System V2 followed by Tecan’s Celero™ EZ DNA-Seq. For control or Ppp1r3c cKO samples, 250 ng of RNA were used for libraries preparation using the TruSeq Stranded mRNA Library Prep protocol (Illumina, #20020594). Libraries were prepared and sequenced by the NIEHS Epigenomics and DNA Sequencing Core as paired-end 151-mers on an Illumina NextSeq 500 instrument.

### RNA-seq analysis

Read pairs were mapped to the mm10 reference genome via STAR v2.5.1b^57^ with parameters “--outSAMattrIHstart 0 --outFilterType BySJout --alignSJoverhangMin 8 -- limitBAMsortRAM 55000000000 --outSAMstrandField intronMotif --outFilterIntronMotifs RemoveNoncanonical”. Counts per gene were determined via featureCounts (Subread v1.5.0-p1)^58^ with parameters “-s2 -Sfr -p” for the Ppp1r3c dataset (reverse-stranded libraries) or with parameters “-s0 -Sfr -p” for the GPI dataset (unstranded libraries). Evaluated gene models were taken from the NCBI RefSeq Curated annotations as downloaded from the UCSC Table Browser on April 21 2021. DESeq2 v1.42.0^59^ was used for principal component analysis and identification of differentially expressed genes (threshold set at FDR 0.05); for the GPI dataset, the individual mouse identifier is included as a variable in the design formula to account for sample pairing. Gene ontology and pathway analysis were analyzed using Enrichr^60–62^ and graphed using R studio.

### Single cell RNA-seq data mining

Dataset for this analysis was taken from Chen et al.^28^, with Dot Plot generated as part of the ShinyCell package (v. 2.1.0)^63^. Differentially expressed genes (DEG) between male and female somatic clusters were determined using the “FindMarkers” function in Seurat (v. 4.3.0)^64^ based on the Wilcoxon Rank Sum test with adjusted p-value < 0.05.

### Germ cell number quantification

Germ cell number in cultured gonads was quantified using FIJI on whole mount samples stained with DDX4 immunofluorescence^55^. For *Ppp1r3c* mice, germ cells were quantified using FIJI and an in house trained CellposeSAM model^65^ by segmenting TRA98 positive cells in subsequent 10 um maximum intensity projection through each whole mount gonad. For Syrosingopine treated gonads the HUCD positive germ cells were counted by hand in FIJI following the same 10um projection paradigm. Data visualization and statistical analysis (two-tailed t-test) and one-way ANOVA was performed using GraphPad Prism.

### Lactate quantification

Lactate concentration was measured using the Lactate-Glo Assay (J5021, Promega). Briefly, single cultured gonads or E12.5 *Ppp1r3c* control or cKO gonadal pairs were dissected from the mesonephros, snap-frozen on dry ice and stored at -80°C until processing. For sample preparation, each sample was homogenized in 50 µl of homogenization buffer (50mM Tris, pH 7.5) and 6.25 µl of inactivation solution (0.6N HCl) using a microtube homogenizer. Immediately after homogenization, 6.25 µl of neutralization solution (1M Tris base Trizma) was added. Protein concentration was determined using 10 µl of the sample with the Qubit Protein Assay (Q33211, ThermoFisher Scientific). For lactate detection, 50 µl of the processed sample was mixed with 50 µl of freshly prepared lactate detection reagent and incubated at room temperature for 60 minutes. A standard curve was generated using serial dilution of lactate, starting at 200 µM, to interpolate lactate concentration from the samples. Luminescence was recorded using the Promega GloMax Discoverer microplate reader (GM3000, Promega). Lactate concentration per sample was determined by interpolation in the standard curve and normalized to protein content. Data visualization and statistical analysis (two-tailed t-test) was performed using GraphPad Prism.

### Syrosingopine treatment in vivo

Five milligrams of Syrosingopine (medchemexpress, CAS: 84-36-6, CAT: HY-N4115) dissolved in corn oil was injected intraperitoneally into C57BL/6 pregnant mothers once on E10.5 and again on E11.5. Control mice were injected with vehicle (DMSO in corn oil) at the same time points. E12.5 gonad-mesonephros complexes were collected and processed for whole mount immunofluorescence.

## Acknowledgments

We are grateful to the NIEHS Molecular Genomics Core, in particular Xin Xu, Erin Smithberger and Molly Cook for the RNA-seq processing and Comparative Medicine Branch for mouse colony maintenance. We would also like to thank Professor Otto Baba for generously providing the anti-glycogen antibody, and the Saltiel Lab at UCSD for the *Pp1r3c* conditional mutant mouse line. This research was supported in part by the Intramural Research Program of the National Institutes of Health (NIH). The contributions of the NIH authors were made as part of their official duties as NIH federal employees, are in compliance with agency policy requirements, and are considered Works of the United States Government. However, the findings and conclusions presented in this paper are those of the authors and do not necessarily reflect the views of the NIH or the U.S. Department of Health and Human Services. This work was supported by the Intramural Research Program (Z01-ES102965 to H.H.-C.Y.) of the NIH, National Institute of Environmental Health Sciences and Duke Light Microscopy Core Facility (NIH Shared Instrumentation grant 1S10RR027867-01). J.S. and B.C. were supported by two grants from the NICHD (1R01HD090050 and R01 HD103064 to BC).

## Author contributions

M.A.E, J.S, B.T, H.H.C.Y and B.C conceived and designed the study. M.A.E and J.S performed the experiments and analyzed the data. S.A.G, Y.Y.C. and M.A.E analyzed the sequencing data. T.M generated the glycogen antibody. M.A.E, J.S, and B.C wrote the manuscript. All authors edited and revised the text.

## Resource Availability

### Lead Contact

Inquiries regarding resources, reagents, and further information should be directed to the corresponding author, Blanche Capel.

### Materials Availability

This study did not generate new unique reagents. Established mouse lines and other reagents are available upon request from the lead contact.

### Data and code availability

Bulk RNA-seq datasets have been deposited at GEO: GSE293016 and GSE293017.

## Supplementary figure legends

**Fig. S1: Glycogen does not accumulate before E12.5.** (A) Immunofluorescence for glycogen (green) and the supporting cell marker SF1 (magenta) in transverse E11.5 male and female gonadal sections. Samples were counterstained with DAPI (blue). (B) Immunofluorescence for SF1, SRY or SOX9 (magenta) and glycogen (green) in transverse E11.5 male gonadal sections. Samples were counterstained with DAPI (blue). (C) Immunofluorescence for glycogen (green) in transverse E11.5 male notochord section. Samples were counterstained with DAPI (blue). (D) Immunofluorescence for PYGB (green) and supporting cell marker SF1 (magenta) in transverse E11.5 male and female gonadal sections. Samples were counterstained with DAPI (blue).

**Fig. S2: Characterization of glycogen metabolism perturbation models.** (A) Darkfield visualization (left) and immunostaining (right) for proliferation (Ki-67, green) and apoptosis (CC3, magenta) in gonadal sections from GPI treated gonads at different concentrations (75 µM, 100 µM and 150 µM) or their respective vehicle solution (DMSO). Samples were counterstained with DAPI (blue). (B) PAS staining on E12.5 control male and female gonads and E12.5 *Ppp1r3c* cKO male gonads. Black dashed box indicates the magnified area.

**Fig. S3: Transcriptomic characterization of glycogen metabolism perturbation models.** (A) Principal component analysis (PCA) plot of the bulk RNA-seq of paired testes cultured with 75 µM GPI or vehicle solution. (B) Volcano plot of differentially expressed genes between 75 µM GPI or vehicle treated testes. Green and cyan dots represent downregulated and upregulated genes, respectively, in the GPI treated testes compared with vehicle controls. (C) Principal component analysis (PCA) plot of the bulk RNA-seq of control and *Ppp1r3c* cKO testes. (D) Volcano plot of differentially expressed genes between the *Ppp1r3c* cKO and the control testes. Blue and red dots represent downregulated and upregulated genes, respectively, in the cKO testis in comparison to the control. Immunofluorescence for the Sertoli cell marker SOX9 (green) and germ cell marker TRA98 (magenta) in E11.5 (E) or E13.5 (F) male control and *Ppp1r3c* cKO gonads. Scale bars=50 μm. (G) Quantification of the total number of germ cells in male control and *Ppp1r3c* cKO testes. Bars represent mean±s.e.m., n≥4. Multiple two-tailed t-test. ns adjpvalue>0.05; * adjpvalue<0.05.

**Fig. S4: Exogenous carbon sources fail to rescue germ cell loss induced by GPI treatment**. 3D projection of DDX4 (germ cell, magenta) whole mount immunostaining in cultured male gonads treated with 75 µM GPI or 75 µM GPI + 25mM sodium lactate (A) or 25mM D-glucose (C). Samples were counterstained with DAPI (blue). Dashed white line delineates the testicular region. Scale bars are 100 μm. Quantification of the total number of germ cells in male gonads cultured with 75 µM GPI or 75 µM GPI + 25mM sodium lactate (B) or 25mM D-glucose (D). Bars represent mean±s.e.m., n=3. Paired two-tailed t-test. ns p>0.05. (E) Quantification of the total number of germ cells in male gonads cultured with vehicle solution, 75 µM GPI, 75 µM GPI + 25mM sodium lactate or 75 µM GPI + 25mM D-glucose. Bars represent mean±s.e.m., n>3. One-way ANOVA. ** p<0.005. (F) 3D printed mold used to generate the agar slabs for single gonad cultures.

### Supplementary table legends

**Table S1: Gonadal single nucleus RNA-seq analysis.** This table contains the differential expression analysis of glycogen synthesis and degradation genes in XX and XY supporting cells at different developmental timepoints.

**Table S2: RNA-seq GPI + pathway analysis**. Bulk RNA-seq data from 75 µM GPI or vehicle treated testes. This table contains QC summary, fragment size plot, fragment size data and the differential expression analysis. This table also contains gene ontology analysis (reactome) for the significantly upregulated and downregulated genes.

**Table S3: Bulk RNA-seq data from E12.5 control and *Ppp1r3c* conditional knockout testes**. This table contains QC summary, fragment size plot, fragment size data and the differential expression analysis. This table also contains gene ontology analysis (biological process) for the significantly upregulated and downregulated genes.

**Table S4: Primary and secondary antibodies used for immunofluorescence.**

