## Supplementary figures and images for "Glycogen metabolism in mouse embryonic Sertoli cells sustains the germ line through the lactate shuttle"

### Supplemental Figures

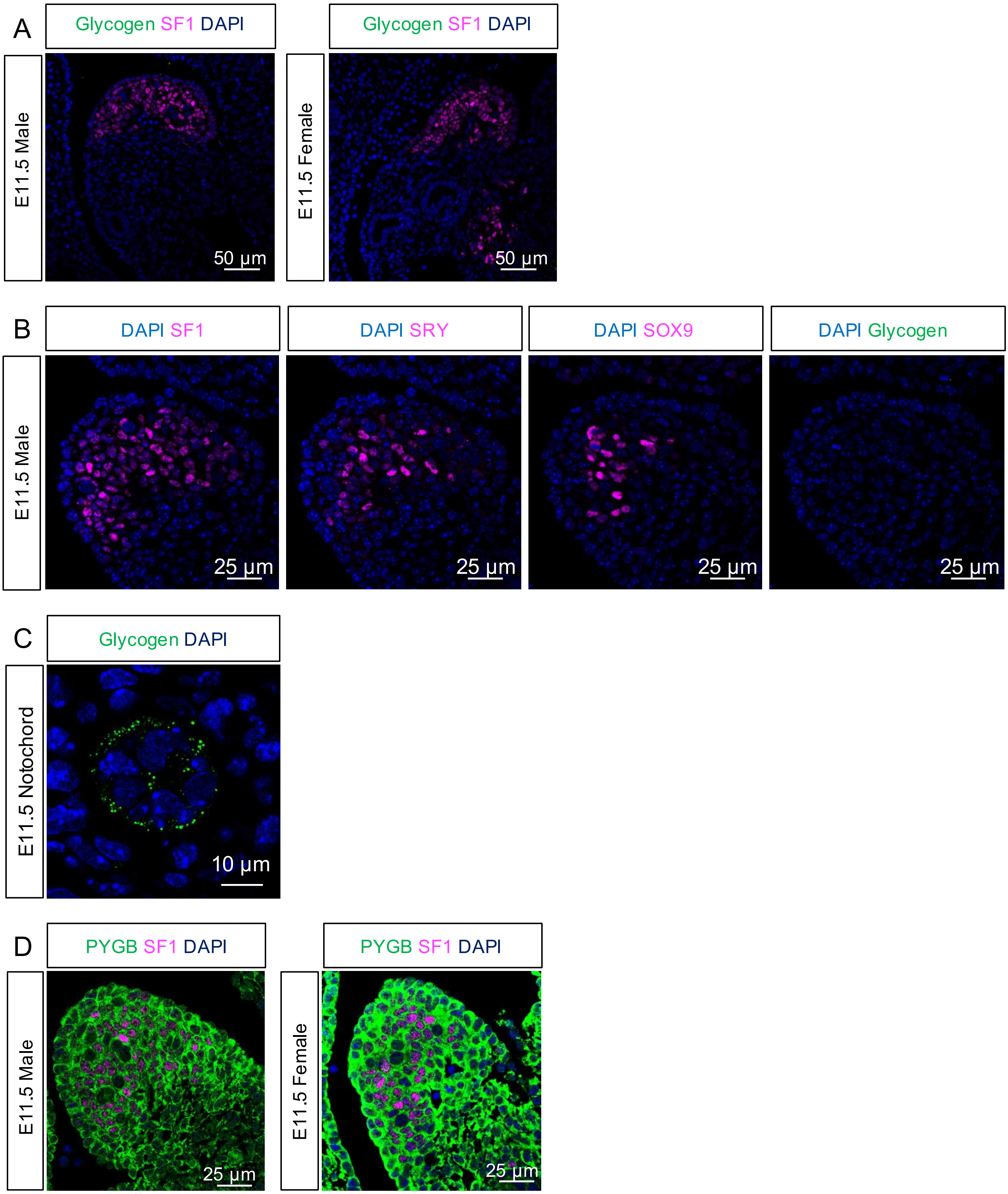

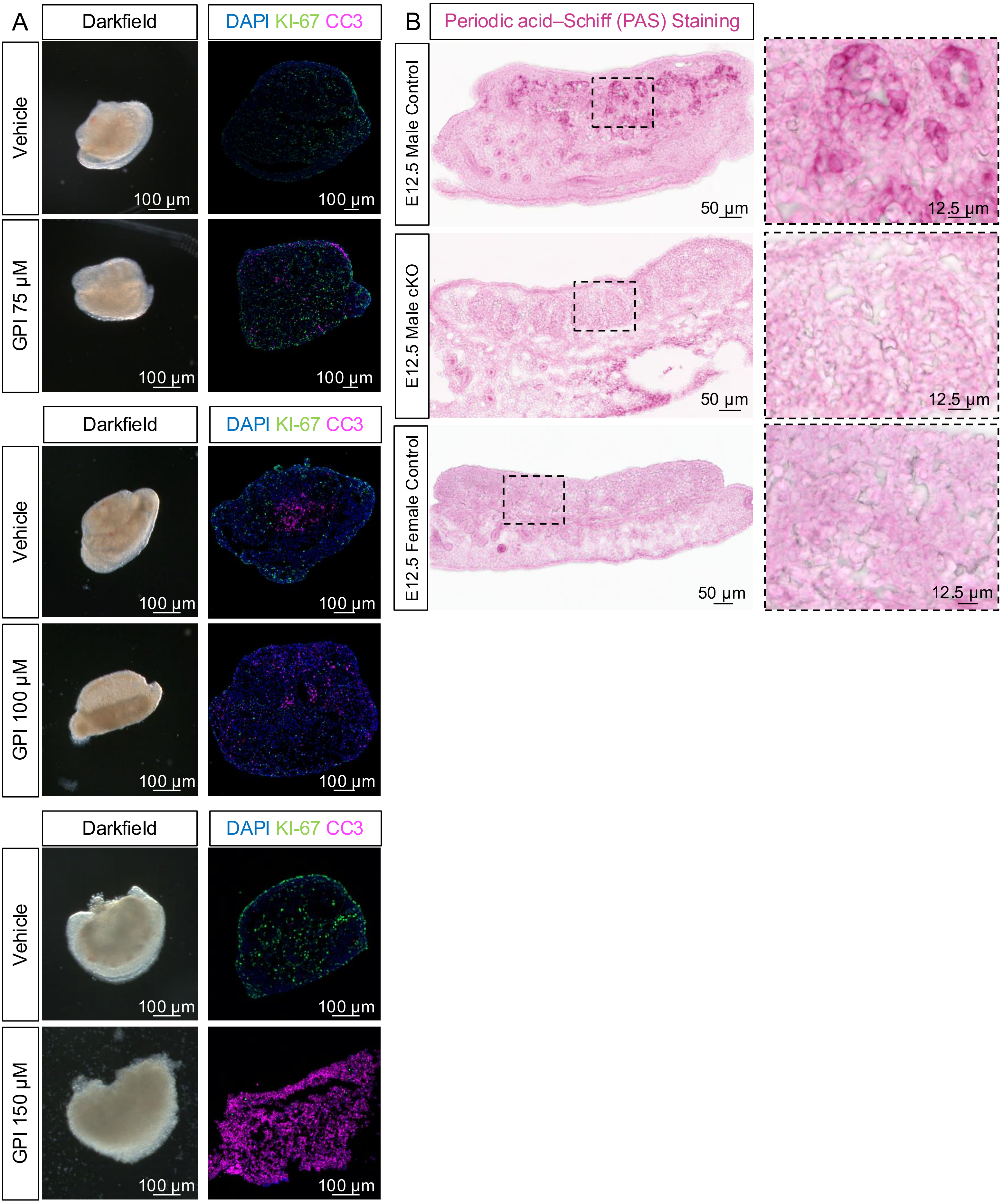

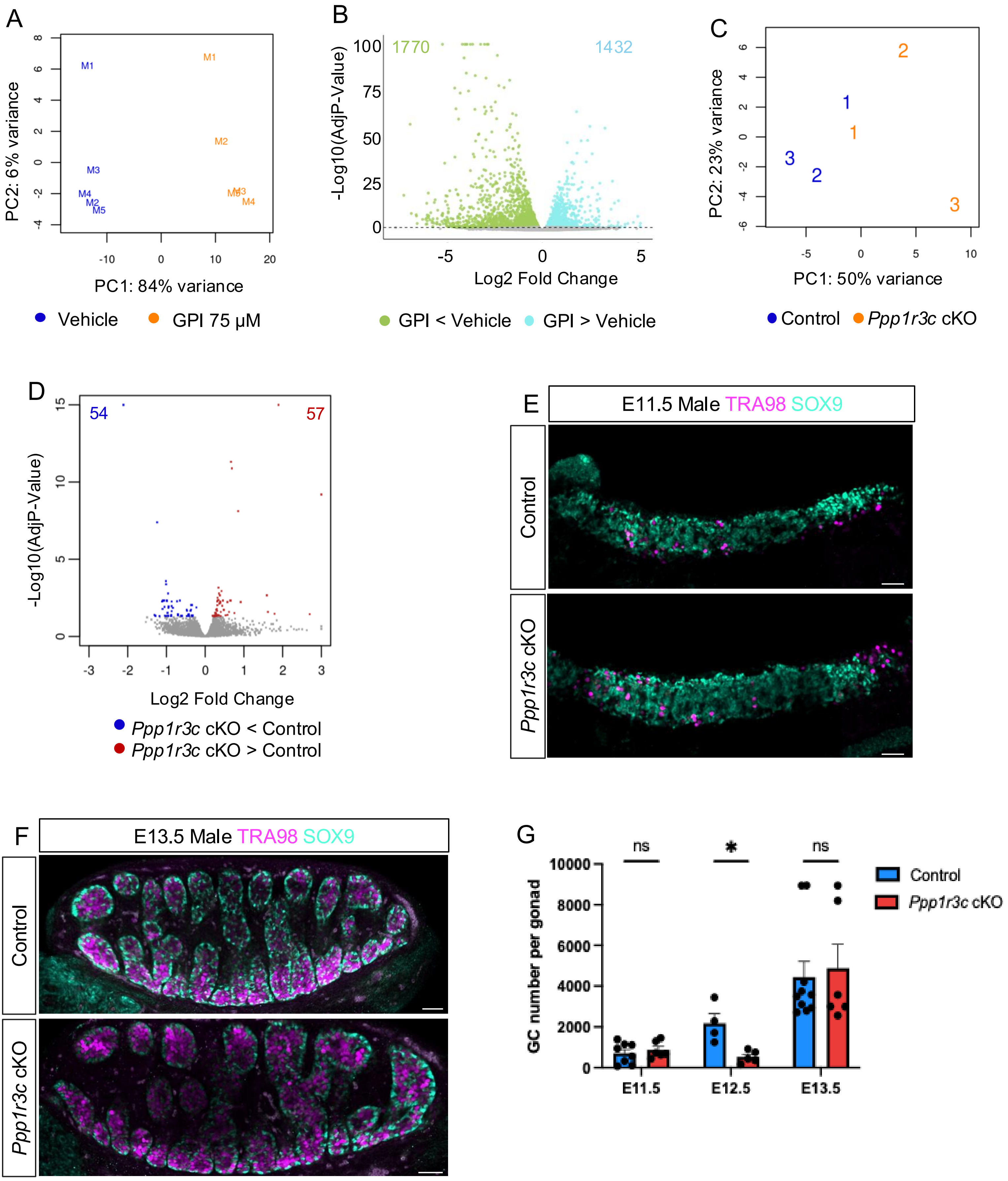

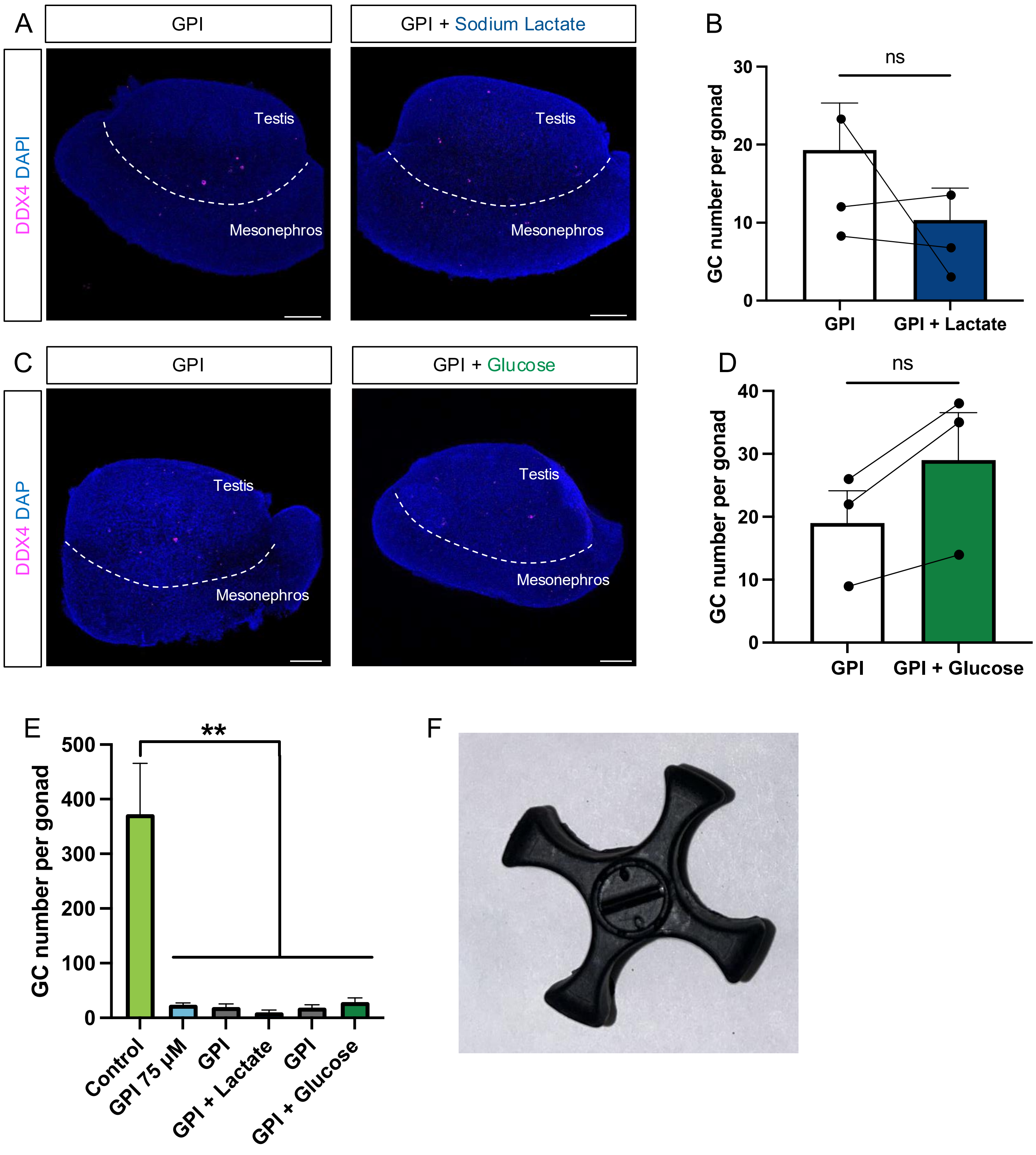
